# Relationship between gut microbiota and circulating metabolites in population-based cohorts

**DOI:** 10.1101/617431

**Authors:** Dina Vojinovic, Djawad Radjabzadeh, Alexander Kurilshikov, Najaf Amin, Cisca Wijmenga, Lude Franke, M. Arfan Ikram, Andre G. Uitterlinden, Alexandra Zhernakova, Jingyaun Fu, Robert Kraaij, Cornelia M. van Duijn

**Author notes:** These authors contributed equally to this work (Dina Vojinovic, Djawad Radjabzadeh, Alexander Kurilshikov). These senior authors contributed equally to this work (Jingyaun Fu, Robert Kraaij, Cornelia M. van Duijn). Correspondence: Prof. Cornelia M. van Duijn, Department of Epidemiology, Erasmus MC, University Medical Center, P.O. Box 2040, 3000 CA, Rotterdam, the Netherlands, Dr. Dina Vojinovic, Department of Epidemiology, Erasmus University Medical Center, P.O. Box 2040, 3000 CA, Rotterdam, the Netherlands.

## Abstract

Gut microbiota has been implicated in major diseases affecting the human population and has also been linked to triglycerides and high-density lipoprotein levels in the circulation. As recent development in metabolomics allows classifying the lipoprotein particles into more details, we aimed to examine the impact of gut microbiota on circulating metabolites measured by Nuclear Magnetic Resonance (^1^H-NMR) technology in 2,309 individuals from the Rotterdam Study and the LifeLines-DEEP cohort in whom gut microbiota was profiled using 16S rRNA gene sequencing. The relationship between gut microbiota and metabolites was assessed by linear regression analysis while adjusting for age, sex, body-mass index, technical covariates, medication use, and multiple testing. Our analysis revealed association of 32 microbial families and genera with very-low-density and high-density subfractions, serum lipid measures, glycolysis-related metabolites, amino acids, and acute phase reaction markers. These observations provide novel insights into the role of microbiota in host metabolism and support the potential of gut microbiota as a target for therapeutic and preventive interventions.

## INTRODUCTION

There is increasing interest in the role of the gut microbiota in the major diseases affecting the human population. For a large part, these associations can be attributed to metabolic and immune signals of the microbiota that enter the circulation.^1^ The gut microbiota has been implicated in obesity and diabetes,^2^ while recently it was also shown that the microbiota is also a substantial driver of circulating lipid levels, including triglycerides and high-density lipoproteins (HDL).^3, 4, 5^ The association with low-density lipoprotein (LDL) cholesterol levels, the major target for treatment of dyslipidemia, or total cholesterol was weaker than the association with triglycerides and HDL.^3, 4^ Recent development in metabolomics allows subclassifying the lipoprotein classes into more detail based on their particle size, composition, and concentration. Various studies further linked the gut microbiota to various amino acids, which have been implicated in diabetes and cardiovascular diseases.^6, 7, 8, 9, 10^

To provide novel insights into the relation of gut microbiota and circulating metabolites, we performed an in-depth study of the metabolome characterized by nuclear magnetic resonance (^1^H-NMR) technology. To obtain sufficient power, we combined the data of two large population-based prospective studies including Rotterdam Study and LifeLines-DEEP cohort, which have a rich amount of data on risk factors and disease.

## RESULTS

Participants from the Rotterdam Study (n = 1,390, mean age 56.9±5.9, 57.5% women) were older compared to the participants from LifeLines-DEEP study (n = 915, mean age 44±13.9, 58.7% women), while sex distribution in the two cohorts was comparable.

The results of association analysis between circulating metabolites (**Supplementary Table 1**) and composition of gut microbiota are illustrated in **Figure 1**. There were 32 microbial families and genera associated with various circulating metabolites after adjusting for age, sex, BMI, medication use and multiple testing (**Figure 1, Supplementary Table 2**). After additional adjustment for smoking and alcohol intake, similar association pattern was observed (**Figure 1B, Supplementary Table 3**). The direction of effect size across the cohorts was generally concordant (Supplementary Table 2 and 3).

**Figure 1.**
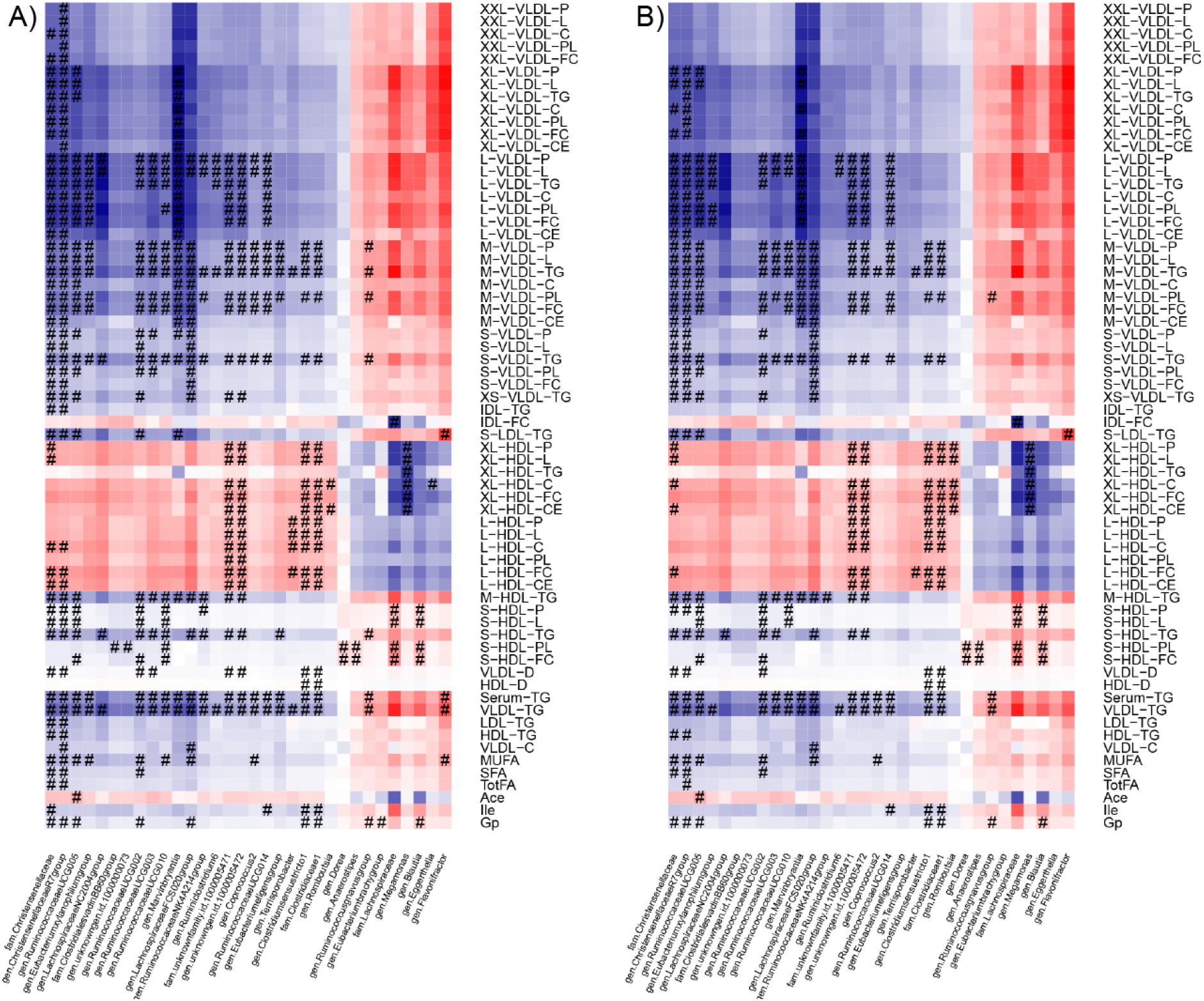
A) Results of association analysis between metabolites and microbial genera and families. Microbial genera and families are shown on *x*-axis whereas metabolites are displayed on *y*-axis. Lipoprotein classes include very low-density lipoprotein particles (VLDL), low-density lipoprotein particles (LDL) and high-density lipoprotein particles (HDL) of very low (XS), low (S), medium (M), large (L), very large (XL) and extremely large (XXL) size. The colors represent effect estimates of the metabolites and microbial taxa after adjustment for age, sex, body-mass index, technical covariates, and medication use. Blue color stands for inverse association. Red color denotes positive associations. Symbols on the plot represent the level of significance with hash denoting Bonferroni significant associations. B) Association between metabolites and microbial genera and families after additional adjustment for smoking and alcohol consumption.

We detected significant associations between 18 microbial families and genera and very low-density lipoprotein (VLDL) particles of various sizes (extra small, small, medium, large, very large, extremely large) and 22 microbial families and genera and HDL particles (small, medium, large, very large) (**Figure 1B, Supplementary Table 3**). There were 13 microbial families and genera associated with both VLDL and HDL particles. For example, family *Christensenellaceae* and genera *Christensenellaceae R7 group, Ruminococcaceae (UCG-005, UCG-003, UCG-002, UCG-010), Marvinbryantia* and *Lachnospiraceae FCS020 group* were found to be associated with VLDL particles of various sizes, small HDL particles and triglycerides in medium HDL particles (**Supplementary Table 3**), whereas family *Clostridiaceae1* and genus *Clostridium sensu stricto 1* were additionally associated with very large and large HDL particles (**Figure 1B**). The strongest association was observed between family *Lachnospiraceae* and free cholesterol in small HDL particles (*p*-value = 1.11×10^-13^) (**Supplementary Table 3**). Of note is that the association pattern of very large and large HDL particles including concentration of particles and its total lipids, cholesterol, free cholesterol, cholesterol esters was opposite compared to the association pattern of small and medium HDL (**Figure 1**).

In addition, there were more targeted associations. For instance, we confirmed previously reported association of genus *Ruminococcus gnavus group* and serum triglycerides and identified novel association with phospholipids in medium VLDL and triglycerides in VLDL (**Figure 1B**).^11^ We also identified a novel association between family *Lachnospiraceae* and genus *Blautia* with small HDL particles (**Figure 1B**) and family *Clostridiaceae1* and genus *Clostridium sensu stricto 1* with both the HDL diameter and VLDL diameter(**Figure 1B**). The VLDL diameter was further associated with family *Christensenellaceae* and genera *Christensenellaceae R7 group* and *Ruminococcaceae UCG-002*.

We also observed association of 15 microbial families and genera with serum triglycerides of which 6 were also associated with fatty acids including monounsaturated (MUFA), saturated (SFA) and total fatty acids (TotFA) (**Figure 1**). These associations followed the same direction of association of VLDL (**Figure 1**).

Beyond the lipoprotein fractions, 8 microbial families and genera were significantly associated with three other metabolites, including the ketone body acetate, amino acid isoleucine, and acute phase reaction marker glycoprotein acetyl (mainly alpha 1) (**Supplementary 3**). Genus *Ruminococcaceae UCG-005* was associated with both acetate and glycoprotein levels, while family *Clostridiaceae1* and genus *Clostridium sensu stricto 1* showed association with both isoleucine and glycoprotein levels. Further, genus *Ruminococcaceae UCG-014* was associated with isoleucine levels, while family *Christensenellaceae* and genera *Christensenellaceae R7 group, Ruminococcus gnavus group*, and *Blautia* showed association with glycoprotein levels.

We next determined whether microbial diversity of gut microbiota was associated with lipoprotein particles or other metabolites (**Figure 2**). When adjusting for multiple testing and age, sex, BMI and medication use, the pattern emerging was that higher microbiome diversity was significantly associated with lower levels of VLDL particles (small, large, medium, very large, extra-large), TotFA, MUFA, and SFA and increased levels of large and extra-large HDL particles and an increased diameter of HDL (**Figure 2**). As to the other metabolites, higher microbiome diversity was significantly associated with lower levels of glycoprotein acetyl, alanine, isoleucine, and lactate (**Figure 2**).

**Figure 2.**
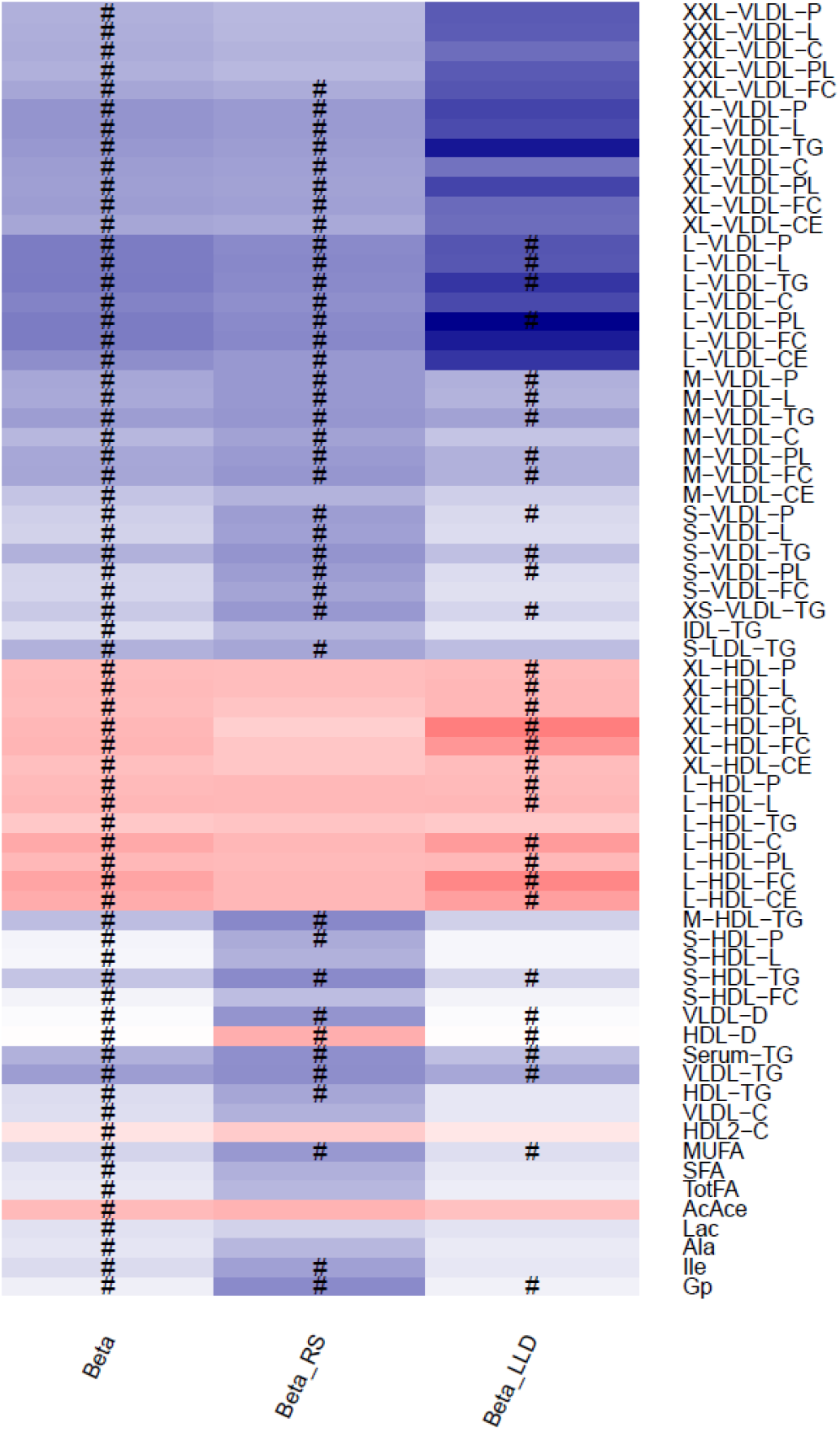
Results of association analysis between metabolites and alpha diversity. Effect estimates from meta-analysis (Beta), and in each of the participating studies are shown (effect estimate in Rotterdam Study - Beta_RS, effect estimate in LifeLines DEEP - Beta_LLD) on *x*-axis, whereas the metabolites are displayed on *y*-axis. Lipoprotein classes include very low-density lipoprotein particles (VLDL), low-density lipoprotein particles (LDL) and high-density lipoprotein particles (HDL) of very low (XS), low (S), medium (M), large (L), very large (XL) and extremely large (XXL) size. The colors represent effect estimates of the metabolites with alpha diversity. Blue color stands for inverse association. Red color denotes positive associations. Symbols on the plot represent level of significance with hash denoting Bonferroni significant associations.

## DISCUSSION

We have examined the impact of gut microbiota on host circulating metabolites in 2,300 individuals from Rotterdam Study and LifeLines-DEEP cohort using ^1^H-NMR technology. We identified associations between the gut microbiota composition and various metabolites including specific VLDL and HDL lipoprotein subfractions, serum lipid measures including triglycerides and fatty acids, glycolysis-related metabolites including acetate and lactate, amino acids including alanine and isoleucine, and acute phase reaction marker including the glycoprotein acetyl independent on age, sex, BMI, and medication use. No associations were found to LDL subfractions except for triglycerides in small LDL and glucose levels measured by ^1^H-NMR.

Our results based on two large population-based studies identified novel associations between the gut microbiota composition and various lipoprotein particles. We observed an inverse association of family *Christensenellaceae* with VLDL particles of various sizes, small HDL particles, and triglycerides in medium HDL (**Figure 1B**). The family *Christensenellaceae* was previously linked to BMI and was associated with the reduced weight gain as reported in the mice study in which germfree mice were inoculated with lean and obese human fecal samples.^12^ Furthermore, the family *Christensenellaceae* was reported to be the most heritable microbial taxon in the study by Goodrich *et al*. independently of the effect of BMI.^12^

Interestingly, the gut microbiota composition showed association with VLDL and HDL particles of various sizes, however weak association has been found for LDL and IDL particles suggesting that gut microbiota affects distinct classes of lipoproteins.^13^ While VLDL particles of various sizes showed the same pattern of association, differences were noticed between large, medium, and small HDL particles suggesting that they are heterogeneous structures.^14^ Small HDL particles are dense, protein-rich, and lipid-poor, whereas large HDL particles are large, lipid-rich particles.^15, 16^ Despite the fact that HDL is consistently associated with a reduced risk of cardiovascular disease, the past decade has seen major controversies on the clinical relevance of HDL interventions. Most trials aiming to increase HDL levels in the aggregate have been unsuccessful and were even stopped because of adverse effects.^17, 18, 19^ The heterogeneity of HDL classes has been long recognized but can now be assessed on a large scale. This compositional heterogeneity of HDL results in functional heterogeneity such that small and large HDL particles are negatively correlated and display inverse relationship with various diseases including cardiovascular disease, as reported previously.^14, 15^ As observed in our study the small HDL particles were driven by genus *Blautia* and family *Lachnospiraceae* and were associated with lower diversity. Indeed the higher levels of small lipoprotein particle concentration have previously been associated with increased risk of stroke as reported in a recently published study of Holmes *et al.*, while the large and extra-large HDL particles that were driven by family *Clostridiacae1*, genus *Clostridiumsensustricto1*, and unknown family and genus and were associated with decreased risk of cardiovascular disease and stroke.^7^ Interestingly, family *Clostridiacae1* was previously inversely correlated with BMI, serum triglycerides and is known to be involved in bile acid metabolism.^11, 20^

Furthermore, we confirmed the association of genus *Ruminococcus gnavus group* and serum triglycerides level,^21^ and additionally reported an association with triglycerides in VLDL particles and phospholipids in medium VLDL. *Ruminococcus gnavus group* was previously associated with low gut microbial richness^22^ and its abundance was higher in patients with atherosclerotic cardiovascular disease.^23^

In addition to circulating lipids and lipoprotein particles, an association was found between gut microbiota and ketone bodies including acetate, amino acids including isoleucine, and acute phase reaction marker including glycoprotein acetyls mainly alpha 1. Circulating levels of acetate were specifically associated with genus *Ruminococcaceae UCG-005*. Acetate is the most common short-chain fatty acid (SCFA) formed by bacterial species in the colon.^24^ SCFA can serve as an energy source, predominately via metabolism in liver.^25, 26^ Previous studies suggested that acetate mediates a microbiota-brain axis and promotes metabolic syndrome.^27^ Circulating levels of isoleucine, an essential branched-chain amino acid, were inversely associated with family *Clostridiaceae1* and genera *Clostridium sensu stricto 1* and *Ruminococcaceae UCG-014* in our sample. Recent studies reported association of circulating levels of isoleucine with diabetes and cardiovascular disease.^8, 28^ Furthermore, isoleucine was reported to be negatively associated with *Christensenellaceae* and positively with *Blautia*.^29^ Even though we observe the same pattern of association between isoleucine and these taxa, the associations did not reach the significance threshold. Also recently, a study focusing on relation of fecal metabolites using mass spectroscopy (Metabolon) and the gut microbiota was published.^6^ Even though the overlap of measured metabolites is limited, amino acids are measured on both platforms. Other amino acids showed a strong association with the gut microbiota but not isoleucine.^6^ However, the concentration of metabolite levels in feces and blood may differ. This is an important field of future research. Lastly, glycoprotein acetyls, a composite marker that integrates protein levels and glycosylation states of the most abundant acute phase proteins in circulation,^30, 31^ was positively associated with genus *Blautia* and *Ruminococcus gnavus group*. Genus *Blautia* is one of the microbial taxa with substantial heritability in twin study, ^12^ and showed strong association with the host genetic determinants which has been associated with BMI and obesity.^32^ Glycoprotein acetyls are associated with other common markers of inflammation.^30, 31^ Circulating level of glycoprotein acetyls have been implicated in inflammatory diseases and cancer, and have been associated with mortality and cardiovascular disease.^7, 8, 33, 34^

The strengths of our study are large sample size, population-based study design, and harmonized analysis in participating studies while correcting for factors such as use of medication and BMI. Merging the data of two large population-based studies allowed us to internally validate the findings. However, our study has also limitations. When exploring circulating molecules, we focused on metabolites measured by Nightingale platform which covers a wide range of circulating compounds.^35^ However, these compounds represent a limited proportion of circulating metabolites, therefore, future studies should focus on metabolites detected by other more detailed techniques.^36^ Further, the gut microbial composition was determined from fecal samples. As gut microbial composition varies throughout the gut with respect to the anatomic location along the gut and at the given site, more complete picture of the gut microbiota could be obtained by getting samples from different locations along the intestines in the future.^13, 37^

Furthermore, when exploring gut microbiota, we focused on 16S rRNA sequencing. Even though broad shifts in community diversity could be captured by 16S rRNA, metagenomics approaches provide better resolution and sensitivity.^38^ Additionally, the cross-sectional nature of our study failed to track changes within each individual. Future studies should focus on collecting stool and blood samples over time for assessment of longitudinal changes. Finally, although our analyses were adjusted for various known confounders, residual confounding remains possible.

To conclude, we found association between gut microbiota composition and various circulating metabolites including lipoprotein subfractions, serum lipid measures, glycolysis-related metabolites, ketone bodies, amino acids, and acute phase reaction markers. Association between gut microbiota and specific lipoprotein subfractions of VLDL and HDL particles provides novel insights into the role of microbiota in influencing host lipid levels. These observations support the potential of gut microbiota as a target for therapeutic and preventive interventions.

## METHODS

### Study population

Our study population included participants from two Dutch population cohorts: Rotterdam Study and LifeLines-DEEP.

The Rotterdam Study is a prospective population-based cohort study that includes participants from the well-defined district of Rotterdam.^39^ The initial cohort included 7,983 persons, aged 55 years or older in 1990 (RS-I).^39^ The cohort was further extended in 2000/2001 by additional 3,011 individuals, aged 55 years and older (RS-II), and in 2006/2008 by adding 3,932 individuals, aged 45 years and older (RS-III).^39^ All participants provided written informed consent. The institutional review board (Medical Ethics Committee) of the Erasmus Medical Center and by the review board of The Netherlands Ministry of Health, Welfare and Sports approved the study.

The LifeLines-DEEP cohort is a sub-cohort of LifeLines study, a prospective population-based cohort study in the north of the Netherlands.^40^ The LifeLines cohort was established in 2006 among participants aged from 6 months to 93 years.^41^ After completion of inclusion in 2013, the cohort includes 165,000 participants.^40^ A subset of approximately 1,500 LifeLines participants aged 18-81 years was included in Lifelines-DEEP.^41^ The LifeLines-DEEP study is approved by the Ethical Committee of the University Medical Center Groningen.^41^ All participants provided written informed consent.

### Metabolite profiling

Quantification of small compounds in fasting plasma samples was performed using ^1^H-NMR technology in both participating studies.^35, 42, 43^ Simultaneous quantification of a wide range of metabolites, including amino acids, glycolysis-related metabolites, ketone bodies, fatty acids, routine lipids and lipoprotein subclasses was done using the Nightingale Health metabolomics platform (Helsinki, Finland). Detailed description of the method can be found elsewhere.^42, 44^ In total there were 145 non-derived metabolite measures quantified in absolute concentration units across the participating studies (**Supplementary Table 1**).

### Gut microbiota profiling

In order to study gut microbiota, fecal samples were collected from participants of Rotterdam Study and LifeLines-DEEP study. 16S rRNA gene sequencing of the V3-V4 (Rotterdam Study) or V4 (LifeLines-DEEP) variable regions was performed using the Illumina MiSeq platform.^41^ A direct classification of 16S sequencing reads using RDP classifier (2.12) and SILVA 16S database (release 128) were used to reconstruct taxonomic composition of studied communities, with binning posterior probability cutoff of 0.8.^45^ All 16S libraries were rarefied to 10,000 reads prior to taxonomy binning. This OTU-independent approach was utilized to decrease domain-dependent bias. The microbial Shannon diversity index was calculated on taxonomic level of genera, using vegan package in R (https://www.r-project.org/). Gut microbiota composition dataset included 1,427 participants from the RS-III cohort that participated in the second examination round at the study center. Metabolite measurements were available for 1,390 Rotterdam Study (RSIII-2) participants. In the LifeLines-DEEP study, gut microbiota composition dataset included 1,186 participants; from them the metabolite measurements were available for 915 participants.

### Statistical analysis

Prior to the analysis, all metabolites were natural logarithmic transformed to reduce skewness of traits distributions. To deal with values under detectable limit (reported as zeros) we added half of the minimum detectable value of the corresponding metabolite prior to transformation. The metabolite measures were then centered and scaled to mean of 0 and standard deviation (SD) of 1. Similarly, to reduce skewness of the distribution of microbial taxa counts, we first added 1 to all taxonomy counts and then performed natural log transformation.

The relationship between metabolites and microbial taxa was assessed by linear regression analysis while adjusting for age, sex, body-mass index (BMI), technical covariates including time in mail and DNA batch effect (only in Rotterdam Study)^46^ and medication use including lipid-lowering medication (395 users in Rotterdam Study and 34 in Lifelines-DEEP), proton-pump inhibitors (258 users in Rotterdam Study and 72 in Lifelines-DEEP), and metformin (67 users in Rotterdam Study and 8 in Lifelines-DEEP). The analyses were further adjusted for smoking and alcohol consumption. Participants using antibiotics were excluded from the analysis. The summary statistics of participating studies were combined using inverse variance-weighted fixed-effect meta-analysis in R (https://www.r-project.org/). In total, 145 overlapping metabolite measures and 455 overlapping microbial taxa were tested for association. As measurements in both metabolomics and gut microbiota datasets are highly correlated, we used a method of Li and Ji to calculate a number of independent tests.^47^ There were 37 independent tests among the metabolite measures and 274 independent tests among microbial taxa. The significance threshold was thus set at 0.05/ (37 × 274) =4.93×10^-6^.

The relationship between metabolites and microbial diversity was also assessed by linear regression analysis while adjusting for age, sex, BMI, technical covariates and medication use (lipid-lowering medication, protein-pump inhibitors, and metformin) in each of the participating studies and summary statistics results were combined using inverse variance-weighted fixed-effect meta-analysis in R (https://www.r-project.org/).

## Data availability

The data is available from the author on request.

## ACKNOWLEDGMENTS

**Rotterdam Study:** The Rotterdam Study is funded by Erasmus Medical Center and Erasmus University, Rotterdam, Netherlands Organization for the Health Research and Development (ZonMw), the Research Institute for Diseases in the Elderly (RIDE), the Ministry of Education, Culture and Science, the Ministry for Health, Welfare and Sports, the European Commission (DG XII), and the Municipality of Rotterdam. Metabolomics measurements were funded by Biobanking and Biomolecular Resources Research Infrastructure (BBMRI)–NL (184.021.007). This work has been performed as part of the CardioVasculair Onderzoek Nederland (CVON 2012-03), the Common mechanisms and pathways in Stroke and Alzheimer’s disease (CoSTREAM) project (www.costream.eu, grant agreement No 667375), Memorabel program (project number 733050814) and U01-AG061359 NIA. Djawad Radjabzadeh was funded by an Erasmus MC mRACE grant “Profiling of the human gut microbiome”. The generation and management of stool microbiome data for the Rotterdam Study (RSIII-2) were executed by the Human Genotyping Facility of the Genetic Laboratory of the Department of Internal Medicine, Erasmus MC, Rotterdam, The Netherlands. We thank Nahid El Faquir and Jolande Verkroost-Van Heemst for their help in sample collection and registration, and Pelle van der Wal, Kamal Arabe, Hedayat Razawy and Karan Singh Asra for their help in DNA isolation and sequencing. Furthermore, we thank drs. Jeroen Raes and Jun Wang (KU Leuven, Belgium) for their guidance in 16S rRNA profiling and dataset generation. The authors are grateful to the study participants, the staff from the Rotterdam Study and the participating general practitioners and pharmacists. **LifeLines DEEP:** LifeLines-DEEP project was funded by the Netherlands Heart Foundation (IN-CONTROL CVON grant 2012-03 to A.Z. and J.F.); by Top Institute Food and Nutrition, Wageningen, The Netherlands (TiFN GH001 to C.W.); by the Netherlands Organization for Scientific Research (NWO) (NWO-VIDI 864.13.013 to J.F., NWO-VIDI 016.178.056 to A.Z., NWO-VIDI 917.14.374 to L.F., NWO Spinoza Prize SPI 92-266 to C.W., and NWO Gravitation Netherlands Organ-on-Chip Initiative (024.003.001) to C.W.); by the European Research Council (ERC) (FP7/2007-2013/ERC Advanced Grant Agreement 2012-322698 to C.W., ERC Starting Grant 715772 to A.Z., and ERC Starting Grant 637640 to L.F.); by the Stiftelsen Kristian Gerhard Jebsen Foundation (Norway) to C.W.; and by the RuG Investment Agenda Grant Personalized Health to C.W. A.Z. also holds a Rosalind Franklin Fellowship from the University of Groningen. The authors thank participants and staff of the LifeLines-DEEP cohort for their collaboration. We thank J. Dekens, M. Platteel, A. Maatman, and J. Arends for management and technical support.

## AUTHOR CONTRIBUTION

Conception and design of the study: DV, DR, AK, JF, RK, CvD. Collection of the data: DR, RK, AK, AZ, LF, CW, JF. Analysis: DV, DR, AK. Interpretation of the data: DV, DR, AK, AZ, NA, RK, JF, CvD. Drafting of the manuscript: DV, CvD. All authors read, revised and approved the final draft.

## COMPETING INTERESTS

None declared.

